# Characterization of the Interaction of Nanobubble Ultrasound Contrast Agents with Human Blood Components

**DOI:** 10.1101/2022.02.11.480110

**Authors:** Michaela B. Cooley, Eric C. Abenojar, Dana Wegierak, Anirban Sen Gupta, Michael C. Kolios, Agata A. Exner

## Abstract

Nanoscale ultrasound contrast agents, or nanobubbles, are being explored in preclinical applications ranging from vascular and cardiac imaging to targeted drug delivery in cancer. These sub-micron particles are approximately 10x smaller than clinically available microbubbles. This allows them to effectively traverse compromised physiological barriers and circulate for extended periods of time. While various aspects of nanobubble behavior have been previously examined, their behavior in human whole blood has not yet been explored. Accordingly, herein we examined, for the first time, the short and long-term effects of blood components on nanobubble acoustic response. We observed differences in the kinetics of backscatter from nanobubble suspensions in whole blood compared to bubbles in phosphate buffered saline (PBS), plasma, or red blood cell solutions (RBCs). Specifically, after introducing nanobubbles to fresh human whole blood, signal enhancement gradually increased by 22.8 ± 13.1% throughout our experiment, with peak intensity reached within 145 seconds. In contrast, nanobubbles in PBS had a stable signal with negligible change in intensity (−1.7 ± 3.2%) over 8 minutes. Under the same conditions, microbubbles made with the same lipid formulation showed a −56.8 ± 6.1% decrease in enhancement. Subsequent confocal, fluorescent, and scanning electron microscopy analysis revealed attachment of the nanobubbles to the surface of RBCs, suggesting that direct interactions, or hitchhiking, of nanobubbles on RBCs in the presence of plasma may be a possible mechanism for the observed effects. This phenomenon could be key to extending nanobubble circulation time and has broad implications in drug delivery, where RBC interaction with nanoparticles could be exploited to improve delivery efficiency.

---

Over the past 30+ years, ultrasound contrast agents (UCAs) have been used in numerous diagnostic and therapeutic applications ranging echocardiography and detection of liver lesions to opening the blood brain barrier and drug delivery.^1–4^ Conventional clinical UCAs are microparticles (known as microbubbles, MBs) with a hydrophobic gas core stabilized by a shell made from phospholipids, proteins, polymers or a combination of these materials. UCAs are visible under ultrasound because of their impedance mismatch, high compressibility of the gas core, and unique nonlinear oscillations in an acoustic field. Because these features are distinct from the surrounding tissue or blood, UCAs can be detected with a high signal-to-noise ratio and imaged with low tissue background.^5,6^ MBs are typically between 1 and 10 μm in diameter and are considered blood pool agents that remain exclusively in the vascular space.^4^ To expand on potential applications of contrast enhanced ultrasound, a new class of UCAs based on nanobubbles (NBs) has been recently developed. NBs are submicron and typically range from 200-500 nm in diameter and have the potential to extravasate due to their smaller size.^4,7–11^ Furthermore, the typical injected *in vivo* concentration of NBs is also 3-5 orders of magnitude higher MBs.^12,13^ The smaller size and higher particle density of NBs facilitates unique applications that can complement those of MBs. For example, NBs can detect biomarkers in the tumor microenvironment or deliver drug beyond the intravascular space.^10,11,14,15^

The properties of the surrounding medium play a considerable role in the behavior of UCAs in an acoustic field.^16–18^ However, in the context of human whole blood, prior experimental work has shown minimal effects of blood cells on the acoustic response of MBs.^19^ This was attributed partially to the size scale of MBs being similar to RBCs. It was concluded that due to the minimal interaction with ultrasound waves at clinical frequencies, blood can be considered Newtonian and homogeneous for modeling contrast agent particle dynamics. In addition to work done with MBs and RBCs, there has been some interest with other blood components, including plasma. Bovine plasma has been used to produce and stabilize albumin-shell MBs.^20^ Because NBs are much smaller and greatly outnumber RBCs (∼10^8^ NBs/mL versus ∼4-6 × 10^6^ RBCs/mL if 5 mLs of NBs are injected into a human), it is critical to examine how their behavior is influenced by fresh human whole blood and blood components. Accordingly, to understand the effects of blood and its components on the acoustic response of NBs, we examined time-dependent acoustic NB behavior in 1) fresh human whole blood, 2) PBS, 3) plasma, and 4) RBCs. Results from this study can be used to gain a better understanding of how the presence of blood cells affects NB imaging properties and potentially be exploited to improve NB performance in diagnostic and therapeutic applications.

## Results

### Nanobubble activity in whole blood

NB suspensions in freshly isolated human whole blood were imaged over 500 seconds in an agarose phantom (solution is contained in a thin inclusion within the agarose, which is approximately the thickness of the imaging plane ∼1 mm, to ensure that the test NB solution was continuously insonated; seen in Figure S1a) using nonlinear contrast mode ultrasound. Ultrasound images were acquired immediately after mixing the NB sample with whole blood, plasma, RBCs, or PBS or after a period of 6, 24, or 48 hours. Mean signal enhancement was measured for each frame of imaging to construct a time intensity curve (TIC), and activity in whole blood was compared to that in PBS. Representative images of NBs in PBS vs whole blood over time and the corresponding TICs can be seen in Figure 1a-b with a representative image in Figure S1b. Representative TICs for NBs in blood imaged immediately or after 48 hours post initial mixing are shown in Figures 1b and 1c. The TICs for all post-mixing time points are shown Figure S2. In all experimental groups, a delayed time to peak enhancement was observed for NBs suspended in whole blood. The average time to reach a steady state enhancement (defined as the time to reach the final enhancement value minus 0.25, which is approximately the standard deviation of the last ten seconds of values) was 145.5 ± 73.4 s. The increase in contrast enhancement when the solution is imaged immediately after mixing was 3.3 ± 1.1 dB and at 48 hours post-mixing, it was 7.5 ± 1.3 dB. After 145 s, the NB signal in whole blood remained stable for the remainder of the experiment (0.8 ± 0.8 dB 145 to 500 s). In contrast, the signal from NBs in PBS showed a stable signal intensity for the entire experiment (−0.4 ± 0.5 dB from 1 to 500 s). However, the signal decay in PBS became more pronounced with increased post-mixing time (Figure 1c), yet no signal decay was observed after any post-mixing time points in whole blood (Figures 1c-d). The delay in time to peak intensity and lack of decay in nonlinear signal from NBs in whole blood was observed regardless of the duration that NBs were exposed to blood before imaging (Figure 1d).

**Fig. 1.**
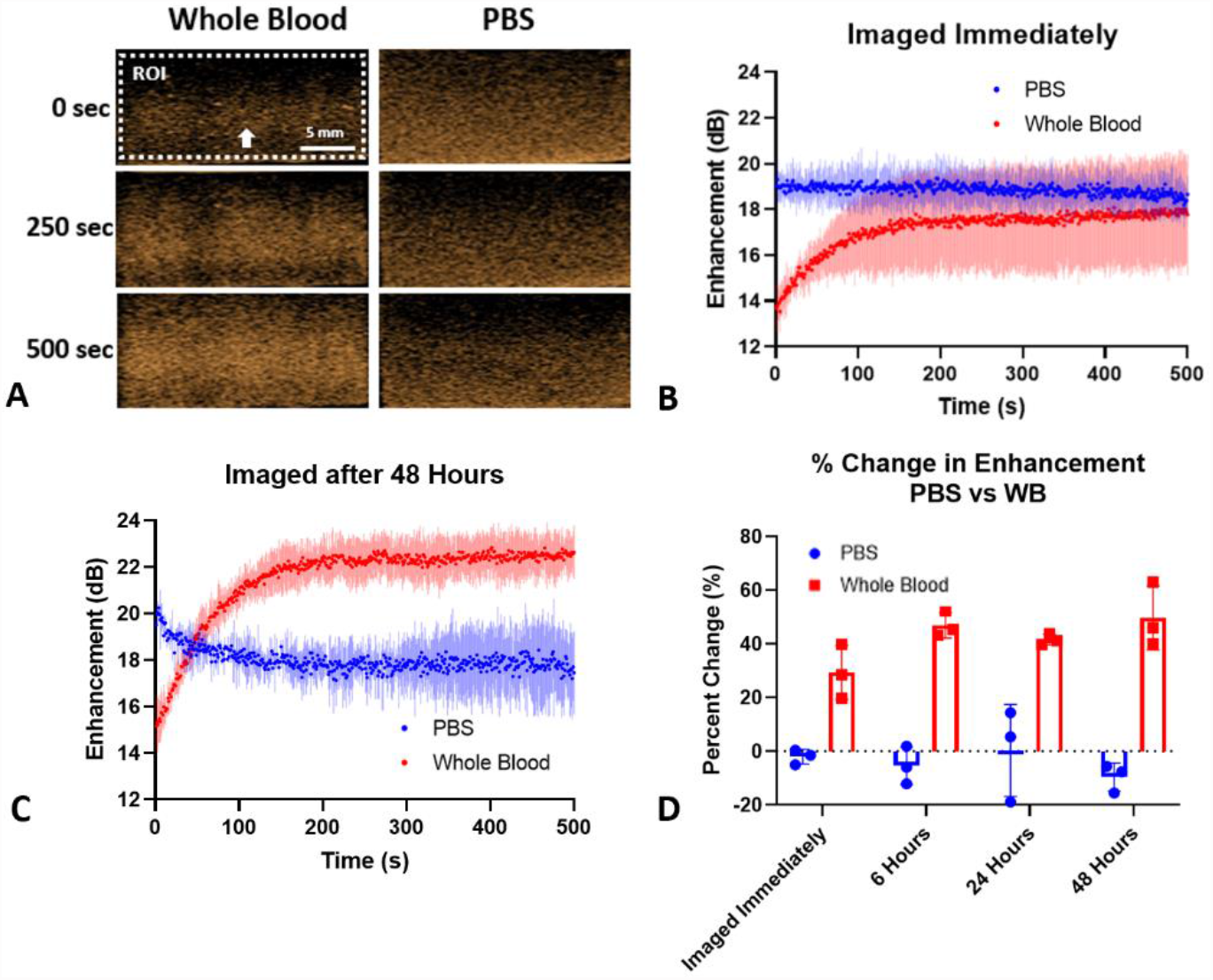
Difference in acoustic activity when the surrounding medium is PBS vs whole blood. (a) Visual representation of the increase in enhancement over time in whole blood. The arrow indicates the direction of the ultrasound beam. (b) When there is no delay in imaging after mixing the NBs and whole blood, the whole blood group reaches the same maximum as the PBS group. There is an increase in enhancement in the first ∼145 seconds in the whole blood group. (c) When there is a delay in imaging post-mixing the solution, the whole blood group continues to increase in enhancement and remain stable, but the PBS group immediately decreases. (d) The percent change in enhancement from the first second to second 500. All whole blood groups show an increase in enhancement. The arrow indicates the direction of the ultrasound beam. n=3 for all groups. Bars represent standard deviation.

### Nanobubble activity in separated blood components

We examined NB activity in separated blood components to explore the potential mechanism behind the delayed time to peak and improved signal stability in whole blood. Here, fresh human whole blood samples were separated into RBCs and plasma, excluding the white blood cell buffy layer. NB acoustic activity under nonlinear imaging was examined in various combinations of these components. Specifically, we examined NB activity in (a) a solution of plasma in PBS, (b) a solution of RBCs in PBS, and (c) a recombination of RBCs and plasma excluding white blood cells. Physiological concentrations of RBCs or plasma were maintained by replacing the missing component with PBS (*e*.*g*. 55% plasma: 45% PBS). The buffy coat, or white blood cell layer, was excluded from all groups. RBCs and plasma were recombined in physiological concentration (55% plasma: 45% RBCs) in the recombination group. NBs were imaged immediately after mixing and after 24 or 48 hours at 25°C. Images were acquired and processed as described above. Representative TICs are shown in Figures 2a-b. All time points can be seen in Figure S3. When the NB solution is imaged immediately after mixing, the NB signal in all groups excluding the recombination group was stable with no significant change in enhancement. As time post initial mixing increases, the plasma and PBS groups showed a decrease in enhancement over time. After 48 hours, the NB signal at 500 s in PBS decreased by −8.4 ± 0.5 dB and the plasma group decreased by −5.0 ± 1.4 dB. In contrast, NB signal in the RBC solution was stable at all time points (*e*.*g*. the change in the enhancement was −0.44 ± 0.74 dB at 48 hours). The percent change in enhancement can be seen in Figure 2c. There was no significant increase in enhancement or delayed time to peak enhancement in any group, at any time point. The peak enhancement never exceeded 19 dB in any group, unlike the previous results in whole blood (Figure 1). The signal from NBs in the RBC suspension was stable in all cases. This suggests that RBCs, particularly in the presence of plasma, likely play a role in stabilizing NBs and decreasing NB dissolution while in the acoustic field. Important to note is that absolute intensity and changes in signal in Figure 2 may differ from Figure 1 due to variability in blood from multiple donors. The data in Figure 1 was conducted with blood from a single donor. The same is true for Figure 2a-b.

**Fig 2.**
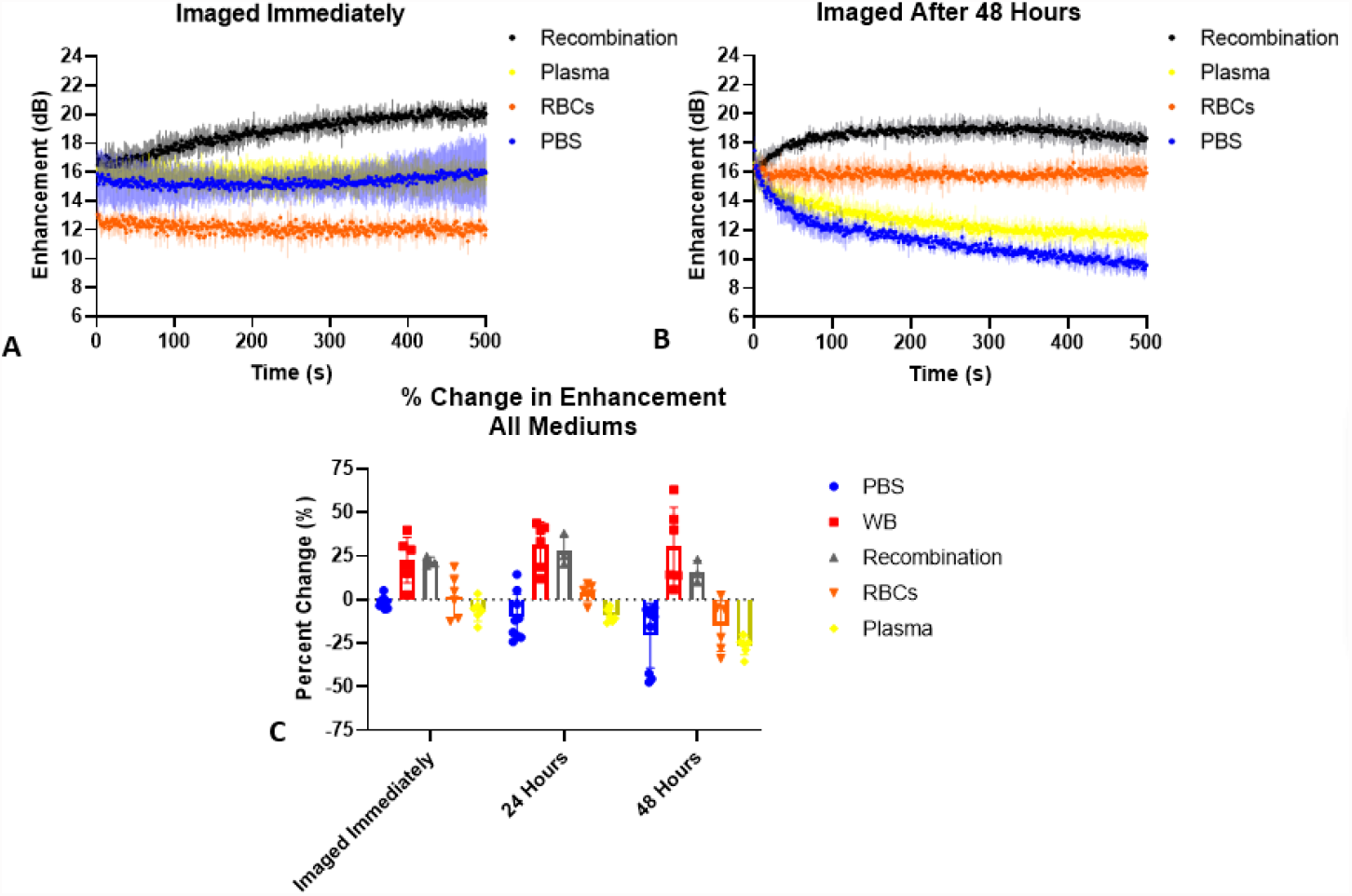
Effect of different blood components on enhancement and stability. (a-b) Nanobubbles in PBS, RBCs, plasma, or recombined plasma and RBC solutions show that PBS and plasma behave similarly, but RBCs have improved stability at later time points post initial mixing. The recombination group shows a delayed time to peak enhancement, similar to the whole blood group in Fig. 1 (n=3). (c) All experimental data (PBS: n=9; whole blood, RBCs, plasma: n=6; recombination: n=3). There is no significant difference between the whole blood and recombination groups from 0-48 hours, but there is a significant difference between PBS, RBCs, and plasma versus whole blood and recombination groups (p < 0.05). Bars represent standard deviation.

To determine if white blood cells played a role in the stability of contrast enhancement and the observed delayed time to peak enhancement, RBCs and plasma were recombined in physiological concentrations (45% RBCs, 55% plasma) and the white blood cell buffy layer was excluded from this group. All experimental data, including this recombination group, were compared to analyze the percent change in enhancement from the first to last second of the experiment (Figure 2c). There was no significant difference between the whole blood and recombination groups imaged immediately or at 24 hours or 48 hours post-mixing, suggesting that white blood cells do not play a critical role in the observed effects. Notably, the slow rise in enhancement was not seen with any group except the RBC-plasma recombination and whole blood groups. This suggests that the slow increase in enhancement is driven by a combination of RBCs and plasma proteins, and is likely a result of direct NB-RBC interactions. To verify that whole blood was not contributing to the change in contrast enhancement, an experimental group with only whole blood (no NBs) was examined. The results can be seen in Figure S4 and show that there is minimal ultrasound signal without NBs.

### Effect of increased insonation on NB activity

To provide additional evidence for the stability of NBs in whole blood and to determine if the delay in time to peak intensity was affected by ultrasonic exposure, the imaging frame rate was increased from 1 fps to 15 fps, and bubbles were imaged for 500 frames (total time: 33 s). The samples were imaged immediately and at 48 hours post-mixing. No delayed time to peak enhancement was observed. Notably, the signal from NBs in whole blood remained stable while the PBS group decayed as the number of frames increased. NBs in PBS showed a −3.9 ± 0.7 dB decrease in enhancement while whole blood had a minimal change in the enhancement of 0.5 ± 0.6 dB. At 48 hours post-mixing, NBs in PBS showed a −2.1 ± 1.2 dB decrease in enhancement while whole blood remained stable with a change in enhancement of −0.2 ± 1.7 dB. The TICs for solutions imaged immediately or at 48 hours post initial mixing of NBs with the solution can be seen in Figures 3a and 3b, respectively.

**Fig 3.**
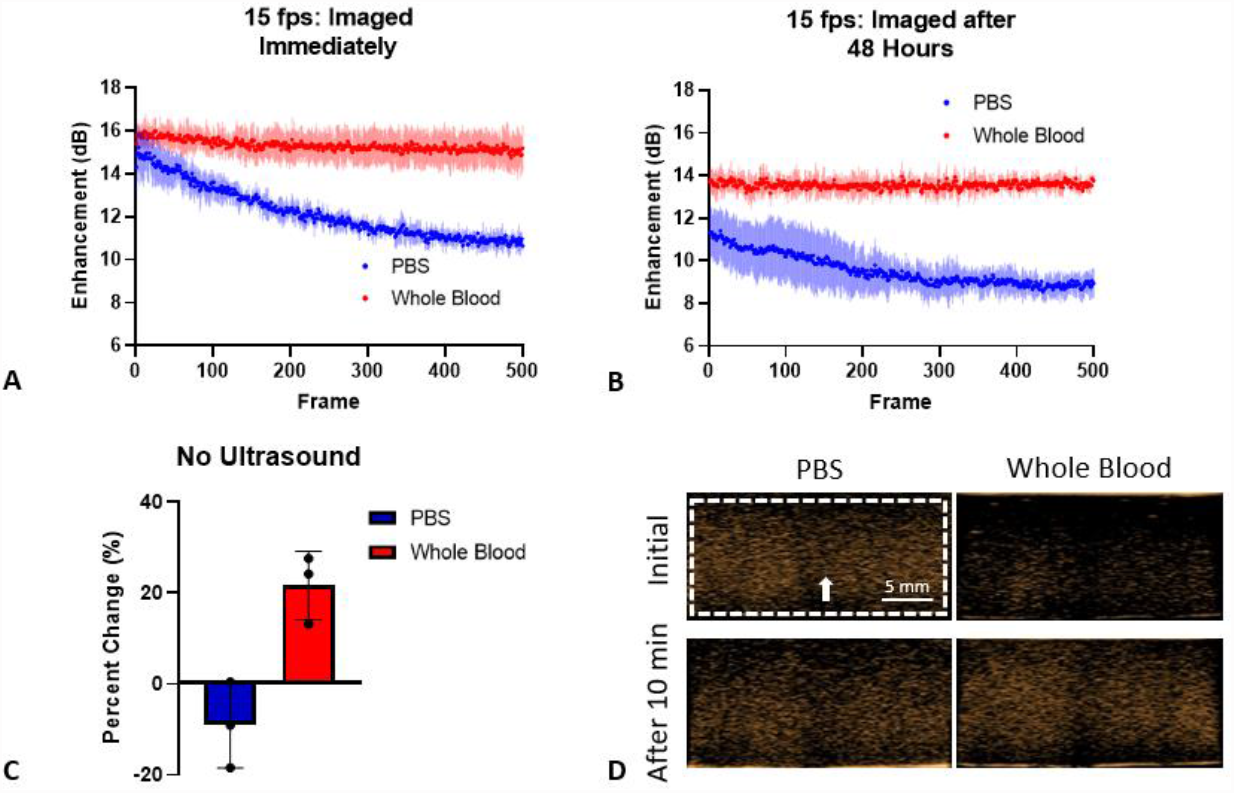
Effect of increasing frame rate or pausing the ultrasound on enhancement. (a-b) Frame rate was increased from 1 fps to 15 fps in PBS and whole blood groups. The PBS groups showed immediate decay in both groups, while the whole blood group remained stable. (c) Percent change in enhancement after the ultrasound was paused for the length of a typical experiment (at 1 fps). The whole blood group showed an increase in enhancement while the PBS groups showed decay. (d) Visual representation of (c). n=3 for all groups. Bars represent standard deviation.

### Temporal effect studies

To examine whether the delayed time to peak enhancement in whole blood was affected by the presence of ultrasound waves, a solution of NBs in whole blood was imaged immediately after it was placed in the agarose phantom inlet and then again after 10 minutes. The ultrasound was paused and no imaging occurred during these 10 minutes. As shown in Figures 3c and 3d, the whole blood group showed a 21.9 ± 7.5% increase in enhancement while the PBS group showed a - 9.0 ± 9.4% decrease in enhancement. This data is consistent with Figure 1a, suggesting that interactions of the acoustic field with NBs in whole blood do not cause the gradual increase in enhancement.

### Effect of varying hematocrit on NB activity

Whole blood was diluted with plasma to examine the direct effect of RBC concentration on NB imaging characteristics. This experiment was important to examine if there is a critical concentration of RBCs required to observe a delay in time to peak intensity. Since plasma, in the presence of RBCs, plays a role in the delayed time to peak enhancement, plasma was not diluted. Results are shown in Figure 4a. 45% (physiological concentration), 34%, and 23% hematocrit (Hct) display similar rises in enhancement (corresponding TICs are seen in Figure 4b). Once the % Hct reaches 11%, there is a slower rise in enhancement and the percent change in enhancement is lower than the less diluted solutions. At 0% Hct, there is no increase in enhancement. There is a significant difference (p < 0.05) between 34% Hct and the following groups: 0% Hct and 11% Hct. There was no significant difference between 45% Hct, 34% Hct, or 23% Hct.

**Fig 4.**
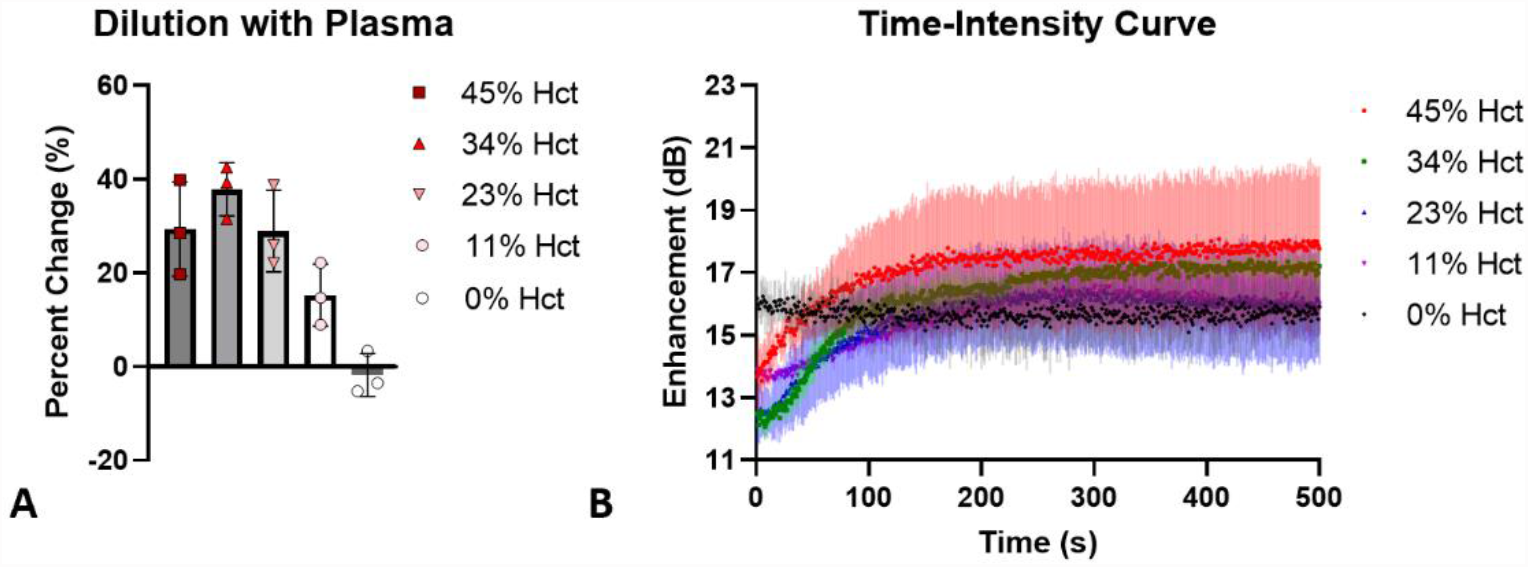
Dilution of whole blood with plasma. (a) As red blood cell concentration (% hct) decreases, the % change in enhancement from the first to last frame of imaging decreases. The observed increase in enhancement becomes less significant as whole blood is diluted with plasma. Bars represent standard deviation. (b) TICs when NBs are in solutions pf varying levels of %Hct. TICs corresponding with Figure 4a. 45%-23% Hct have a similar rise in enhancement while 11% Hct has a more leveled off rise in enhancement. 0% Hct has no increase in enhancement and is stable over the length of the experiment.

### MB behavior in whole blood

To examine how bubble size and shell material affect imaging properties in whole blood, experiments were conducted with commercial MBs (Lumason), and house-made MBs that have the same shell material as the NBs, isolated from the same lipid solutions. MBs were tested in whole blood and PBS using the same setup as for the NBs. Results are shown in Figure 5. Notably, no delayed time to peak enhancement was observed in any MB group. However, the signal decay in whole blood was slower than in PBS for both Lumason and the house-made MBs. The slower decay may indicate a stabilizing effect of whole blood or blood components on MBs and decreased gas diffusion out of the bubble. House-made MBs were also imaged 48 hours post initial mixing in PBS and whole blood (Figure 5c). Similar to the NB experiments (Figure 1), MBs imaged 48 hours post initial mixing with whole blood showed a slower signal decay than MBs mixed with PBS at the same time point. Figure 5d shows a direct comparison of NB and MB TICs in whole blood, imaged immediately after mixing. A difference in signal decay between MBs and NBs can be seen, suggesting that the potential interaction between RBCs and bubbles is size-dependent and more likely to occur with smaller, submicron bubbles.

**Fig 5.**
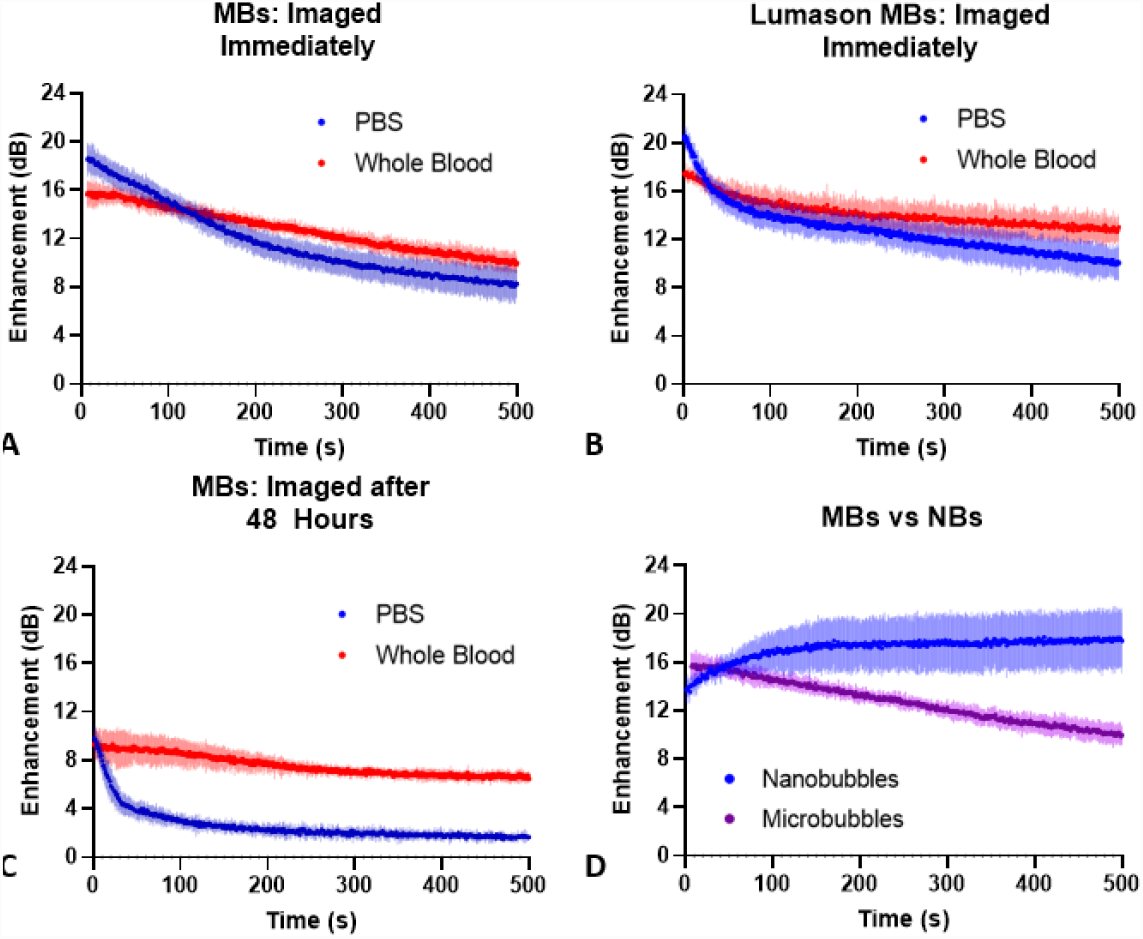
Microbubble stability in whole blood vs PBS. (a, c) MBs with the same lipid shell formation as the NBs were studied imaged immediately after mixing or 48 hours post initial mixing. (b) Lumason MBs were studied and imaged immediately after mixing. Neither house-made microbubbles nor Lumason showed an increase in enhancement. When imaged 48 hours post-mixing, the PBS group showed immediate decay. For all three experiments, the whole blood group had a slower decay than the PBS group. (d) Direct comparison of house-made MB and NB experiments (imaged immediately after mixing). n=3 for all groups.

### Autocorrelation studies

NB-generated speckle can provide important information on randomness of NB movement. This information can provide evidence supporting NB association with RBCs in whole blood, since any non-random association should result in a longer decorrelation time. The less random the NB motion, the longer the decorrelation time. NBs in whole blood showed a significantly longer decorrelation time (23.3±17.7 s) compared to NBs in PBS (1.3±0.5 s), plasma (1.0±0.0 s) or RBCs (1.7±0.5 s) (Figure 6a-b). This suggests that movement of NBs in whole blood is less random than in RBCs, plasma, or PBS. NB attachment to the surface of the RBCs may be causing diminished NB motion in whole blood. This effect was not seen with MBs (Figure 6c); no significant difference was seen between MBs in whole blood (1.7±0.6 s) and PBS (1.7±1.1 s). We hypothesize that this effect is due to adhesion of NBs on RBCs, in the presence of plasma. RBC interactions could be reducing the degrees of freedom for NB movement through attachment and/or acting as physical barriers, decreasing the random Brownian-like motion and slowing the NB signal decay.

**Fig 6.**
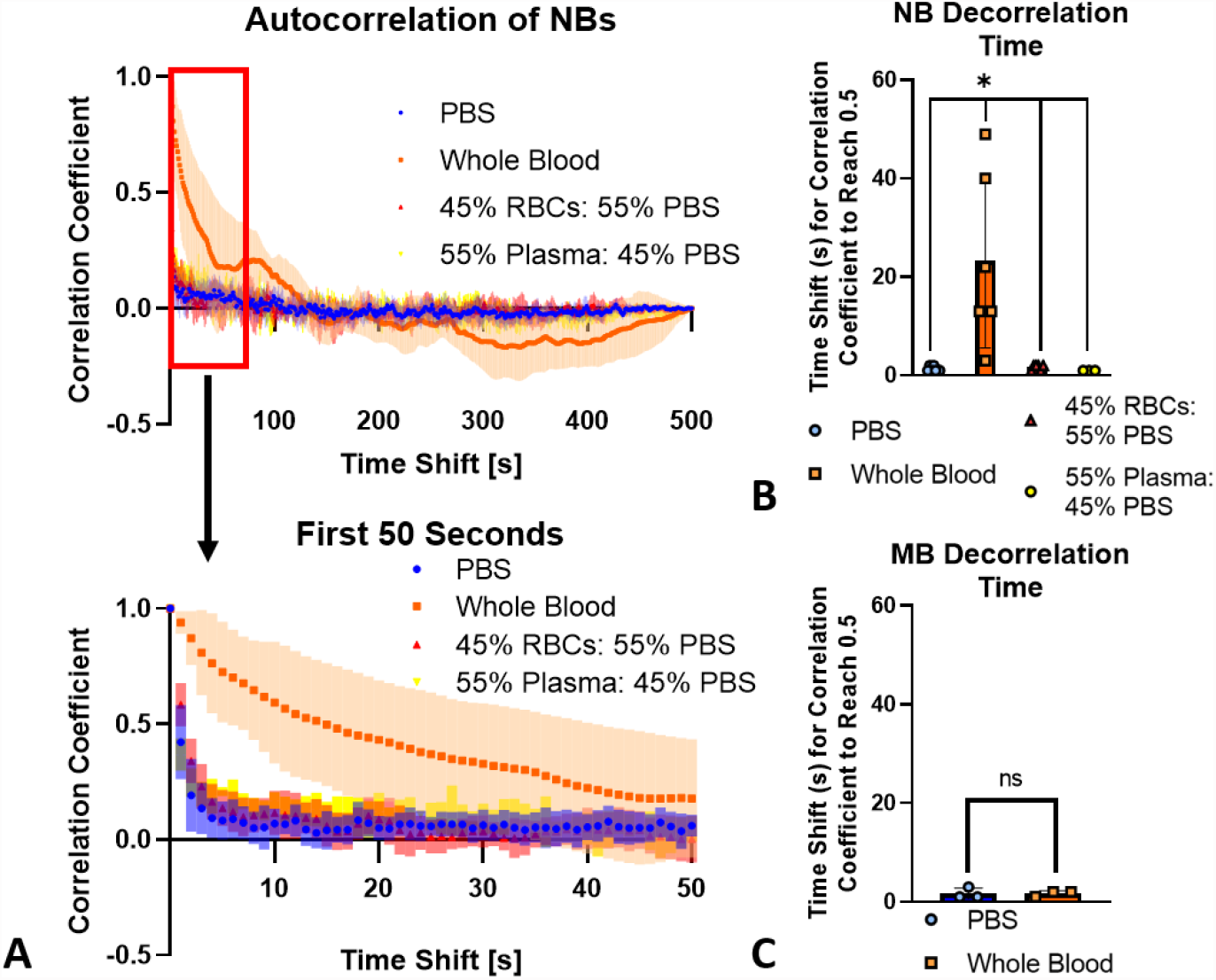
Autocorrelation curves and decorrelation time for NBs and MBs in different solutions. (a) Autocorrelation curves of NBs in whole blood (n=6), PBS (n=7), plasma (n=3) or RBCs (n=6). The first 50 s are zoomed in to clearly highlight the slow decline in correlation of the whole blood group. (b) Decorrelation time (decorrelation is defined as a correlation coefficient of 0.5). p<0.05. (c) MB decorrelation time. The ROI for each image was taken at the focus of the ultrasound and was ∼0.1 × 0.1 mm. Bars represent standard deviation.

### Microscopy

Fluorescent microscopy was performed within 30 minutes of placing NB-whole blood solutions onto a slide. Both fluorescent and brightfield images at 100x indicate that nanobubbles localize around RBCs (Figure 7a). To get a more detailed image of the NB location on individual cells, confocal microscopy was performed four hours after placing the NB-whole blood solution onto a slide to allow for increased interactions between NBs and RBCs. As shown in Figure 7b, NBs can be seen in solution, and some NBs can be seen adhering to the surface of RBCs. Because the NB concentration is roughly three orders of magnitude higher than that of RBCs (4.07 × 10^9^ NBs/mL vs ∼4-6 × 10^6^ cells/mL), it is expected that free NBs will be more readily visualized. Nonetheless, confocal microscopy results show that NBs can interact directly with RBCs.

**Fig 7.**
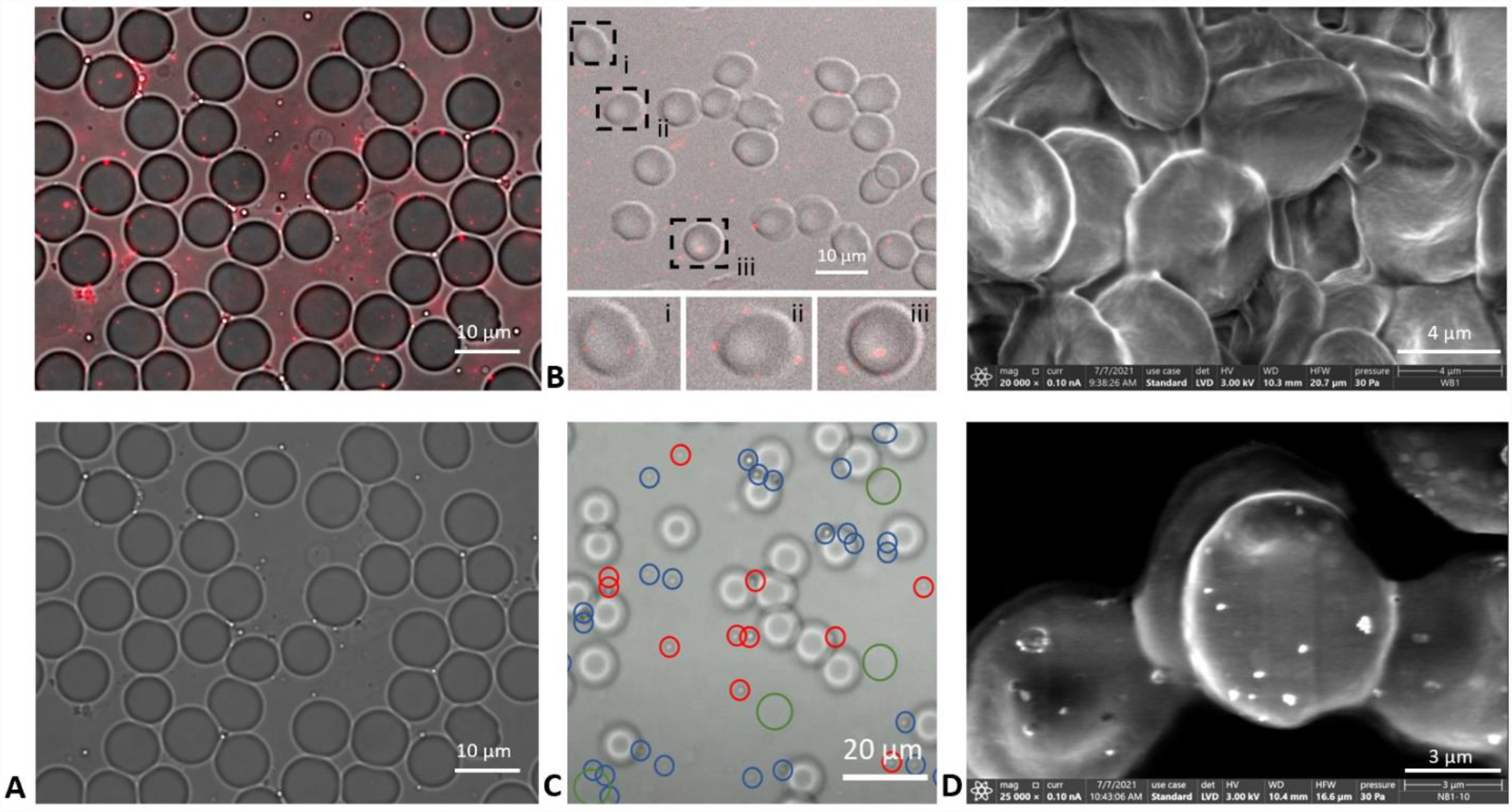
Microscopy of NBs in whole blood. Figure 7.(a) 100x images taken with an inverted fluorescent microscope with a Cy5.5 channel overlaid with brightfield (top) a brightfield alone (bottom). (b) Confocal microscopy image with three RBCs highlighted. There is significant fluorescence around the RBC membrane. (c) 63x brightfield microscopy image taken from a video with the plane of focus directed at the NBs (RBCs appear out of focus). Red circles indicate NBs that moved location from t=0 to t=4 s and blue circles indicate NBs that did not move location. Larger green circles show RBCs that are out of plane. (d) SEM images with whole blood without NBs at 20,000x (top) and with NBs at 25,000x (bottom).

To investigate if the fluorescent and confocal observations were due to adhesion, a 4 s video was acquired at 63x with brightfield microscopy to show unfixed NB and cell movement over time (Figures 7 and S5). NBs circled in blue did not move in the 4 s period, but NBs circled in red did (Figure 7c). Additional still images and the video can be viewed in Figure S5. The immobilized NBs tend to accumulate around the rim of RBCs. It should be noted that some RBCs are out of the plane of imaging, examples of which can be seen with green circles. Some NBs are surrounding out of plane RBCs. The video was taken out of plane with RBCs to make the NBs more visible. The corresponding in plane image can be seen in Figure S5a.

In accordance with previous work discussing nanoparticle hitchhiking, scanning electron microscopy (SEM) was performed (Figure 7d)^21–26^. Unlike other hitchhiking nanoparticles, NBs are gas-filled, highly compressible structures. Thus, on SEM, it is likely that the NBs have collapsed or deflated because the sample must be dried before it can be imaged. According to past work with this formulation of NBs, a majority (greater than 10x) of NBs are buoyant, or gas-filled, as opposed to non-buoyant nanoparticles^13^. When whole blood without NBs (Figure 7d top) was imaged and compared to whole blood with NBs (Figure 7d bottom), there was a stark difference in results. Only when NBs were present did the RBCs appear to have particles on their surface. These particles are approximately the size of the NBs.

## Discussion

The acoustic activity of shell-stabilized gas bubbles depends on many factors, including the bubble compressibility, the shell viscoelastic properties and properties of the surrounding medium. Bubble response in an acoustic field, in turn, directly affects the imaging properties (e.g. how long a signal lasts and how bright an image appears) of the contrast agent. This study examined how fresh human whole blood and blood components influenced bubble response under the nonlinear contrast mode of a clinical ultrasound scanner. Human blood is different from bovine and rodent blood through characteristics such as RBC deformability, membrane rigidity, lifespan, and size.^27,28^ Therefore, it is important to conduct these studies using human blood.

To our knowledge, the stability and nonlinear contrast imaging properties of NBs in human whole blood versus the frequently studied phosphate-buffered saline (PBS), or less frequently studies animal blood, have not been previously explored. Likewise, there have been no studies examining the interactions between RBCs and NBs. This includes the potential physical interaction between NBs and cells and their effect on their acoustic activity. Studies examining MBs and RBCs are more common than NBs. Many studies have focused on how MBs have a similar velocity profile to RBCs in flow, which makes them an ideal model to study blood movement and identify flow abnormalities.^29–32^

There has been significant recent interest in the interactions of other (non-bubble) nanoparticles with RBCs. Prior work has shown that, depending on surface charge and size, some nanoparticles can “hitchhike,” or adsorb through non-covalent interactions, onto the RBC’s surface, which improves their circulation time because it delays physiological clearance.^21–23,25,26,33^ Specifically, the non-covalent adsorption of nanoparticles onto the RBC surface has been shown to reduce uptake into reticuloendothelial system organs by over twofold and this increased accumulation in the target organ (*e*.*g*. lungs) by up to 8.9-fold.^22^ The presence of serum, the size of the particles, and the polymer used to fabricate the particles have all been shown to significantly impact the number of particles adhering to RBCs.^25^ Extended circulation time and increased tumor uptake can be essential for improved molecular imaging target detection and for drug delivery, and in the case of UCAs, potentially longer imaging times.

In this study, the acoustic activity of NB solutions in whole blood versus PBS showed two key differences: 1) a delayed time to peak enhancement in, approximately, the first 145 seconds of imaging at 1 fps and 2) improved stability in an acoustic field at different frame rates and following various time periods post initial mixing. This study showed that there is a delay of approximately 145 s for NBs in whole blood to reach peak intensity and this delay only occurred when both plasma and RBCs were present in the sample. The delay does not seem to be caused by the presence of the acoustic field (Figure 3c). When an autocorrelation analysis was conducted, NBs in the presence of whole blood had a significantly longer decorrelation time compared to NBs in plasma or RBCs alone. This suggests that it was not only the physical presence of cells restricting NB movement since the presence of plasma was required for a longer decorrelation time (i.e. less random movement of the NB signal under ultrasound). Microscopy studies showed that this could be due, in part, to NB associations with the RBC surface. These ultrasound TIC and autocorrelation findings were not seen with MBs.

The delay in time to peak intensity effect did not occur when NBs were in solutions of PBS or blood components (i.e. plasma and RBCs) (Figures 1 and 2). However, when plasma and RBCs were recombined, a delayed time to peak intensity and improved stability were both observed (Figure 2). Since the recombination group excluded white blood cells, we hypothesize that both effects are dependent primarily on the presence of RBCs and plasma, but not white blood cells, in the surrounding solution. NB movement was more restricted in solutions of whole blood when compared to PBS, RBCs, or plasma, indicated by the longer decorrelation time (Figure 6). Based on the experimental observations, we hypothesize that the delayed time to peak intensity and improved stability are driven, in part, by physical interactions between NBs and RBCs in the presence of plasma proteins.

There is likely a critical concentration of RBCs required for the observed effects to occur, and this is shown in the experiment that diluted whole blood with plasma (Figure 4). Studies have shown that the percent hematocrit (Hct) makes a significant difference in the backscattering coefficient in Doppler imaging.^34,35^ The percent Hct may also play an integral role in contrast harmonic imaging with ultrasound contrast agents. The experiments here show that as the % Hct decreases, there is less of an increase in enhancement (i.e. delay in time to peak enhancement). At 0% Hct, there was no percent change in enhancement over time, meaning that there is no delay in time to peak enhancement. This experiment also supports the hypothesis that RBCs are a significant factor contributing to the delayed time to peak and increased stability seen with NBs in whole blood because in this experiment, only the RBC concentration changed; plasma was not diluted. However, it must be noted that as whole blood is diluted, the viscosity of the solution also decreased. Therefore, viscosity could play a role in the greater ultrasound signal stability (lower signal decay) observed.

The effect of fluid viscosity on MB oscillations has been shown to decrease MB oscillation amplitude.^16,36^ Further, MB fragmentation decreases at a higher surrounding fluid viscosity and MBs in more viscous fluid show greater stable cavitation at higher pressures.^18^ Similar results were observed in our study where NBs in whole blood were significantly more stable (echogenic for a longer time period) than NBs in PBS, a less viscous fluid (Figure 5). The viscosity of plasma is about 1.8 times that of water, while the viscosity of whole blood is 3-4 times the viscosity of water.^37^ If NBs are experiencing a higher degree of stable cavitation, as seen in the aforementioned studies, this may partially explain why the maximum echogenicity of NBs in whole blood reached values as high as 22.6 ± 1.2 dB, compared to 17.6 ± 1.3 dB in PBS at the same time point (Figure S2).

While viscosity may play a role in some of our observations, the experimental groups with whole blood (collected in sodium citrate tubes) and RBCs in PBS should have similar viscosities because of their equivalent cellular content (approximately 45% Hct). Whole blood will have a slightly higher viscosity due to the presence of plasma. In these groups, there was a delayed time to peak enhancement when observing solutions of NBs in RBCs without plasma, although there was improved temporal stability (Figure 2). This corresponds to other studies in literature where increased viscosity of the surrounding fluid decreased MB fragmentation and increased stable cavitation.^16,18^ Future experiments could further clarify the role of viscosity and cellular content by using viscous solutions or polymers such as Dextran to differentiate the roles of plasma proteins and viscosity.

Another potential process influencing the observed effects is RBC aggregation and settling.^38,39^ The blood samples are obtained in sodium citrate tubes, which would prevent the sample from clotting and should reduce RBC aggregation, regardless of NB association. Even in the unlikely event that RBC aggregation occurred, RBCs are linear scatterers of ultrasound that are not expected to contribute to the contrast-enhanced ultrasound signals detected in this work. In terms of sedimentation, normal erythrocyte sedimentation rate (ESR) is ≤ 15 or ≤ 20 mm/hr in men and women younger than 50 years old respectively.^40^ It is thus unlikely that there would be significant RBC settling over 8.5 minutes (the length of the experiment) that was not directly visualized in the field of view.

The second observed effect, increased stability of NB signal over time, was shown in experiments at 1 fps and 15 fps (Figure 3). *In vivo*, a higher frame rate is often desirable to improve the temporal resolution of images and typically, there is increased bubble destruction as the frame rate increases, causing ultrasound signal to decrease.^41^ However, in this case there was no decay in enhancement when NBs were imaged in whole blood. In PBS, there was a significant decay of 3-4 dB over 500 images. There was no delay in time to peak enhancement in whole blood at 15 fps. This may be because the increased number of ultrasound pulses per unit time prevented the solution from forming as many NB-RBC interactions compared to imaging at 1 fps, or there may be more NB destruction at 15 fps.

In this study, we showed distinct effects visualized with ultrasound when NBs were in the presence of plasma and RBCs. Prior work on nanoparticle-RBC interactions examined the role of plasma on binding.^25,26^ In general, there appears to be less binding to the RBC surface in the presence of plasma with other nanoparticles. Multiple groups have also tested the effect of zeta potential on RBC interactions and showed that zeta potentials close to zero did not induce toxicity with RBCs and positively charged particles had prolonged *in vivo* circulation time.^25,42,43^ However, the group that found a prolonged circulation time with positively particles did not test any particles with a zeta potential that was negative and as close to zero as our particles. While this was not studied directly herein, it is possible that for NBs, association with RBCs only occurs in the presence of plasma due to the near-neutral surface charge. We have previously found that our nanobubbles have a zeta potential of −2.15 ± 1.78 mV.^44^ It is also possible that plasma has a direct effect on the particle shell properties, which may influence their interactions with RBCs. This is something that will be investigated in future work.

Confocal, fluorescent, brightfield, and scanning electron microscopy provide further evidence on the interactions between RBCs and NBs (Figure 7). Localization of fluorescent NBs around the RBC membrane is observed, indicating that there are NBs on the RBC surface. It should be noted that there is significant background fluorescence, but this was expected because the number of NBs/mL is three orders of magnitude greater than the number of RBCs/mL. Additional evidence for NB adhesion to the RBC surface is seen in Figures 7c and S5 where a video was taken over 4 seconds and NB movement was identified. A majority of NBs that appear fixed were localized to the RBC surface while the free-floating NBs experienced more movement. This video is essential in providing evidence that NBs are adhered to the RBCs and are not just being imaged in the same location as them. SEM indicates that there are NBs present on the RBC surface. Whole blood samples imaged without NBs did not have anything resembling particles on their surface (Figure 7d). SEM is one of the standard imaging modalities for nanoparticle-RBC adsorption studies^21–26^.

An important question in NB research that has yet to be fully explored is why NBs have a longer lifespan *in vivo* than MBs. The possible role of interactions between NBs with blood cells has not been studied. In these experiments, we found that MBs (∼ 1 μm in diameter) of the same formulation as our NBs (∼ 275 nm in diameter) did not exhibit a delay in time to peak enhancement, and the US signal began to decay immediately after imaging began (Figure 5). It should be noted that the concentration of NBs and MBs were ∼4.07 × 10^9^ NBs/mL and ∼1.18 × 10^6^ MBs/mL, respectively, and this could be a confounding factor. The NB and MB solution concentrations were chosen so that the samples had the same starting enhancement in PBS (∼ 20 dB) for direct comparison with no signal attenuation. Note that the number of MBs injected *in vivo* is also significantly smaller than the number of NBs injected. The commercially available MB, Lumason, was also tested, and it also showed a rapid decay in signal enhancement, similar to our MB. Concentration plays a significant role in contrast stability over time and may play a more significant role in MBs than NBs.^45^

While the MB sample did not show an increase in signal enhancement, the signal decay in whole blood was slower than in PBS. Signal decay over 500 seconds of imaging at 1 fps in house-made MBs was 55.4 ± 4.2% in PBS and 36.2 ± 2.5% in whole blood; in Lumason, decay was 50.8 ± 5.4% in PBS and 25.4 ± 5.1% in whole blood (Figure 5). Thus, while the MBs may not be interacting with RBCs in the same way or to the same extent as NBs, whole blood may still stabilize the MBs. This is not necessarily dependent on bubble shell formulation because both our MBs and Lumason exhibited a decay in signal over time.

Bubble size may play a significant role in the delayed time to peak intensity and improved signal stability observed with NBs because when bubbles of the same formulation were tested, only NBs, and not MBs, showed a delay in time to peak enhancement (Figure 5d). Because NBs are significantly smaller than MBs and RBCs, they would be more likely to have multiple NBs congregate around the RBC surface. As NBs adhere to the RBCs, they would be less likely to be taken up by the reticuloendothelial system and in particular, by Kupffer cells, which are an important limiting factor in nanoparticle drug delivery to tumors.^46^

There are limitations to this study. The experiments were conducted in a static environment and at room temperature. Additional studies at body temperature could yield new insights into the observed effects. Likewise, it is not yet known if the RBC-NB associations will occur *in vivo*. Future experiments will examine these effects in animal models. It is likely that if interactions do occur *in vivo*, they will do so in environments with low shear and slow flow, such as in capillaries and venules, to allow for time for associations to occur. *In vivo* interactions could result in distinct kinetics of NBs in small vessels as well as in tumors where the vasculature is complex.

## Conclusion

We examined, for the first time, the acoustic response of lipid-shelled NBs in fresh human whole blood. Experimental results from these studies show that there are distinct differences between NB acoustic response in whole blood versus PBS, RBCs, and plasma. Similar effects were not seen with MBs. These experiments provide an explanation for why some NBs have a longer lifespan *in vivo* than MBs. These results are broadly applicable to improving the circulation time of all nanoparticles, but uniquely focus on lipid-shelled, highly deformable particles. Future work should explore the possibility of NB-blood interactions in a dynamic flow environment and *in vivo* to determine if they can be harnessed for imaging and drug delivery purposes.

## Methods

### Materials

The phospholipids 1,2-dipalmitoyl-sn-glycero-3-phosphate (DPPA) and 1,2-dipalmitoyl-sn-glycero-3-phosphoethanolamine (DPPE) were obtained from Corden Pharma (Liestal, Switzerland); 1,2-dibehenoyl-sn-glycero-3-phosphocholine (DBPC) from Avanti Polar Lipids Inc. (Pelham, AL); and 1,2-distearoyl-snglycero-3-phosphoethanolamine-N-[methoxy(polyethylene glycol)-2000] (DSPE-mPEG 2000) from Laysan Lipids (Arab, AL). Propylene glycol was purchased from Sigma Aldrich (Milwaukee, WI), glycerol from Acros Organics, 1x phosphate buffered saline (PBS) from Gibco (Life Technologies), and C_3_F_8_ perfluorocarbon (PFC) gas from AirGas (Cleveland, OH). Glutaraldehyde was purchased from Alfa Aesar (Ward Hill, MA).

### Whole blood preparation

Anonymized fresh human whole blood was obtained from healthy donors with informed consent via venipuncture through a protocol approved by the Institutional Review Board (IRB) at University Hospitals of Cleveland (IRB Number: 09-90-195) or Case Western Reserve University (IRB Number: STUDY20191092). All blood was collected in sodium citrate containing tubes and used the day of collection unless otherwise noted. For experiments in plasma or RBCs in PBS, whole blood was separated by centrifugation (500g for 10 minutes) and the platelet rich plasma and RBC layers were reconstituted with PBS to reach physiological concentrations (55% plasma and 45% RBCs). The plasma group was composed of 45% PBS and 55% plasma and the RBC group was 55% PBS and 45% RBCs by volume. The recombination group was composed 45% RBCs and 55% plasma, recombined after centrifugal separation, excluding the white blood cell layer. For experiments with varying percent hematocrit (Hct), whole blood was diluted with plasma from the same donor to yield 34, 23, and 11% Hct.

### Preparation of NBs

NBs were prepared via mechanical amalgamation as previously described by de Leon *et al*. ^44^ Briefly, 6.1 mg of DBPC, 2 mg of DPPE, 1 mg of DPPA, and 1 mg mPEG-DSPE 2000 were dissolved in 0.1 mL of propylene glycol via sonication and heating at 80°C. A solution of 0.1 mL of glycerol and 0.8 mL of PBS was heated at 80°C and added to the lipid solution. The solution was sonicated at room temperature for 10 minutes, 1 mL was sealed in a 3 mL headspace vial with a rubber stopper, and sealed with an aluminum cap. The gas inside the vial was replaced with C_3_F_8_ gas by purging the vial with 10 mL of gas. NBs were activated via agitation in a VialMix mechanical shaker (Bristol-Meyers Squibb Medical Imaging, Inc., N. Billerica, MA) and isolated by centrifugation (50g for 5 minutes). NBs had a concentration of ∼4.07 × 10^11^ bubbles/mL (measured prior to introduction to blood) obtained via resonance mass measurement (Archimedes, Malvern Panalytical Inc., Westborough, MA).^44^

### Preparation of MBs

MBs with the same shell composition as the NBs were prepared as recently described by Abenojar *et al*. ^45^ Briefly, the MBs (1 μm diameter) were isolated by diluting the bubbles obtained after VialMix with 10% vol/vol glycerol/propylene glycol in PBS, and centrifuging at 300 g for 10 min after which the infranatant (liquid suspension) was discarded. Next, the MB cake was re-dispersed in PBS and centrifuged at different speeds (30, 70, 160, and 270 g) for 1 minute, after which the cake was discarded and the infranatant collected. The desired MB sample was then obtained following centrifugation at 300 g for 10 min wherein the infranatant was discarded and the cake was re-dispersed in a solution of 10% vol/vol glycerol/propylene glycol in PBS. The sample was then transferred to a 3 mL headspace vial, capped with a rubber septum, and sealed with an aluminum cap. The vial was then flushed with PFC gas and used immediately for imaging. MB shell composition was previously shown to be identical to that of the NBs and had a concentration of ∼1.18 × 10^6^ bubbles/mL (measured prior to introduction to blood).^45^ This concentration was chosen so that NBs and MBs had the same initial enhancement. It would have been difficult, if not impossible, to conduct this study with the same concentration of NBs and MBs. Too high of a MB concentration will cause attenuation and too small of a NB concentration will have minimal ultrasound signal. In addition to house-made MBs, the commercially available MB Lumason (Bracco) was used. Lumason was activated and frozen to prevent decay of bubbles over time. Lumason was thawed in a vial immersed in an ice bath to decrease the decay rate of bubbles between experiments. The concentration of frozen buoyant Lumason bubbles (1.14 ± 0.6 × 10^8^ bubbles/mL) was comparable to never frozen bubbles (3.21 ± 1.1 × 10^7^) (Figure S6).

### Ultrasound imaging experimental setup

Bubble imaging was carried out in agarose phantoms with an inlet for the experimental solution. The phantoms were formulated with 1.5 wt% agarose in Millipore water and were prepared according to de Leon *et al*. ^44^ The phantom and ultrasound setup can be seen in Figure S1a. A separate agarose phantom was used for each experiment.

Experimental solutions included: 1) whole blood, 2) PBS, 3) RBCs with no plasma (45% Hct), 4) plasma with no RBCs, 5) recombined plasma and RBCs (excluding white blood cells), 6) 75% whole blood: 25% plasma (34% Hct), 7) 50% whole blood: 50% plasma (23% Hct), and 8) 25% whole blood: 75% plasma (11% Hct). All experimental groups had the same concentration of NBs (∼4.07 × 10^9^ bubbles/mL based on population measurements from de Leon *et al*.).^44^ Note that if a solution is denoted as “whole blood”, the sample contains unaltered blood (e.g. white blood cells are present). Solutions were imaged at 25°C immediately after adding NBs to the solution and after time periods post initial mixing of 6, 24 and 48 hours. In groups that were not imaged immediately, the sample was mixed with a pipette prior to imaging to prevent settling and separation of sample components. Groups allowed to interact for additional hours post initial mixing remained at 4°C until they were imaged at 25°C, consistent with literature focused on nanoparticle-RBC interactions.^26,43^ Immediately after the solutions were combined or after 24 or 48 hours, the solution was imaged at 25°C in an agarose phantom (inlet volume: 220 mm^3^) with a Toshiba clinical ultrasound imaging system (AplioXG SSA-790A) and a 12 MHz center frequency linear array transducer (PLT-1204BT). Ultrasound images were acquired in contrast harmonic imaging mode with 0.1 MI, 65 dB dynamic range, and 70 dB gain. The frame rate for all experiments was 1 frame per second (fps) unless otherwise noted. 500 frames were acquired for all experiments.

### Temporal effect studies

To investigate if the experimental results are impacted by continuous ultrasound exposure, studies were conducted where the ultrasound was turned off. This helped to determine if ultrasound directly impacts the temporal TIC effects observed. Images were taken immediately after the solution was added to the agarose phantom. The ultrasound was then paused for 10 minutes, and the solution was imaged again. The contrast enhancement from the initial images was compared to the images taken after 10 minutes.

### Analysis of ultrasound data

All experiments were carried out in triplicate unless otherwise noted. Ultrasound images were analyzed by taking the entire phantom as the area of interest, measuring the signal intensity in the region of interest (ROI) (22 mm × 10 mm), and normalizing the data by subtracting the area outside of the phantom as background. An example of an image can be seen in Figure S1b. All experiments were conducted at least 3 times to account for differences in phantom alignment with the ultrasound transducer. The mean enhancement was analyzed statistically between groups by unpaired Student’s t-tests. The difference between the initial intensity and final intensity of each experiment was analyzed statistically by paired t-tests. A p-value of < 0.05 was considered significant.

### Autocorrelation analysis

Ultrasound signal decorrelation time was used to measure the random Brownian-like motion of bubbles in solution exposed to ultrasound. The analysis was conducted using house-made NB and MB samples. NB solutions included PBS, whole blood, 45% RBCs: 55% PBS, and 55% plasma: 45% PBS. MB control experiments were carried out using whole blood and PBS. Autocorrelation coefficient values were calculated using the smallest ROI possible (0.1 mm × 0.1 mm) using MATLAB. Decorrelation time was defined as the time shift required to reach a correlation coefficient of 0.5.

### Microscopy

NB solutions in whole blood were imaged using confocal, fluorescent, scanning electron, and light microscopy. DSPE was conjugated with Cy5.5 and incorporated into the NB membrane through agitation. Cy5.5 conjugated to DSPE lipid was prepared by reacting Cyanine5.5 maleimide (Lumiprobe, Maryland, USA) and DSPE PEG Thiol (DSPE-PEG-SH (2k) (Nanocs, New York, USA)) and was dissolved in chloroform in a 1:2 mol ratio. The reaction was allowed to proceed for at least 2 hours at room temperature and stored at −20 °C afterwards. A Leica TCS SP8 STED confocal microscope from the CWRU (Case Western Reserve University) Light Microscopy Imaging Core Facilities was used in both brightfield and fluorescent channels and overlaid. Images were taken after a time period of 4 hours to decrease the movement of the cells and bubbles. During this time delay, the sealed and prepared slides were enclosed in aluminum foil and left at 4°C.

Cy5.5-conjugated NBs in whole blood were imaged within 30 minutes (imaged immediately) of activation and preparation on a glass slide with a cover slip using a Zeiss Axio-Observer inverted fluorescent microscope at 63x and 100x. Images were taken in brightfield and with a Cy5.5 fluorescent channel.

Scanning electron micrographs were collected on the ThermoFisher Apreo 2S (CWRU, Case School of Engineering SCSAM Center, NSF MRI Award: 2018167) in low vacuum mode. Samples were prepared according to McGregor *et al*.^47^ Briefly, a human whole blood sample was drawn the day before imaging and fixed with 4% glutaraldehyde for 1 hour. Samples were either fixed with whole blood or whole blood with nanobubbles at a ∼4.07 × 10^10^ bubbles/mL concentration. The sample was transferred to PBS until 1 hour before imaging. 5 μLs of each sample were dried on a silicon wafer and then imaged.

## Supporting information

Video for Supplemental Figure 6d

## Acknowledgements

This research was supported by the Hematopoietic Biorepository and Cellular Therapy Shared Resource of the Case Comprehensive Cancer Center (P30CA043703); the NIH grants T32 GM007250 and T32 HL134622; and the National Institute of Biomedical Imaging and Bioengineering (R01EB025741 and R01EB028144). The authors would like to acknowledge the support of the staff and use of the ThermoFisher Apreo 2S (NSF MRI Award: 2018167) in the CWRU, Case School of Engineering SCSAM Center for the collection of the scanning electron micrographs. The authors would also like to acknowledge the support of R. Lee from the Case Western Reserve University Light Microscopy Imaging Core.

## Supporting Information Available

Lumason size and concentration after freezing the sample; experimental setup; all experimental groups for enhancement experiments; enhancement of whole blood with no NBs; light microscopy and video of NBs in whole blood

