## Supplemental Figures for "Characterization of the Interaction of Nanobubble Ultrasound Contrast Agents with Human Blood Components"

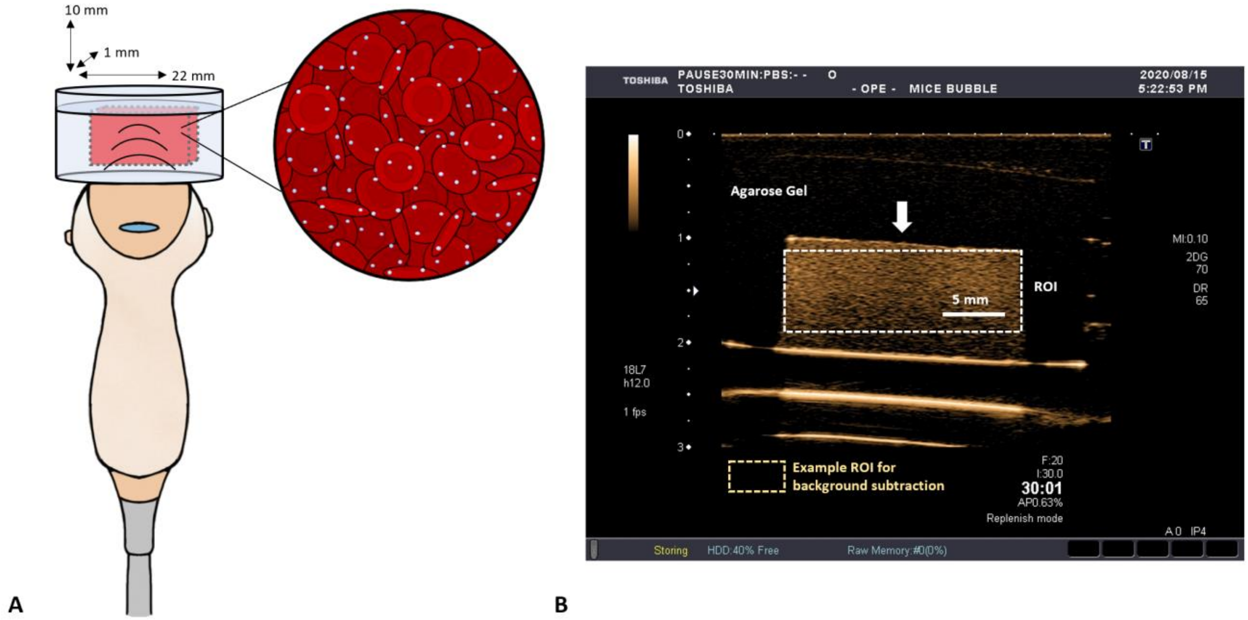


**Supplemental Figure 1. Experimental setup.** (a) Agarose phantom setup with inlet for experimental solution. Nanobubbles can be seen in blue on the expanded image. (b) Example of an ultrasound image seen with this setup. The ROI used for analysis can be seen with the white rectangle and an example of a background ROI can be seen with the yellow rectangle. The arrow represents the direction of the ultrasound beam, which is inverted for convenience in other figures.

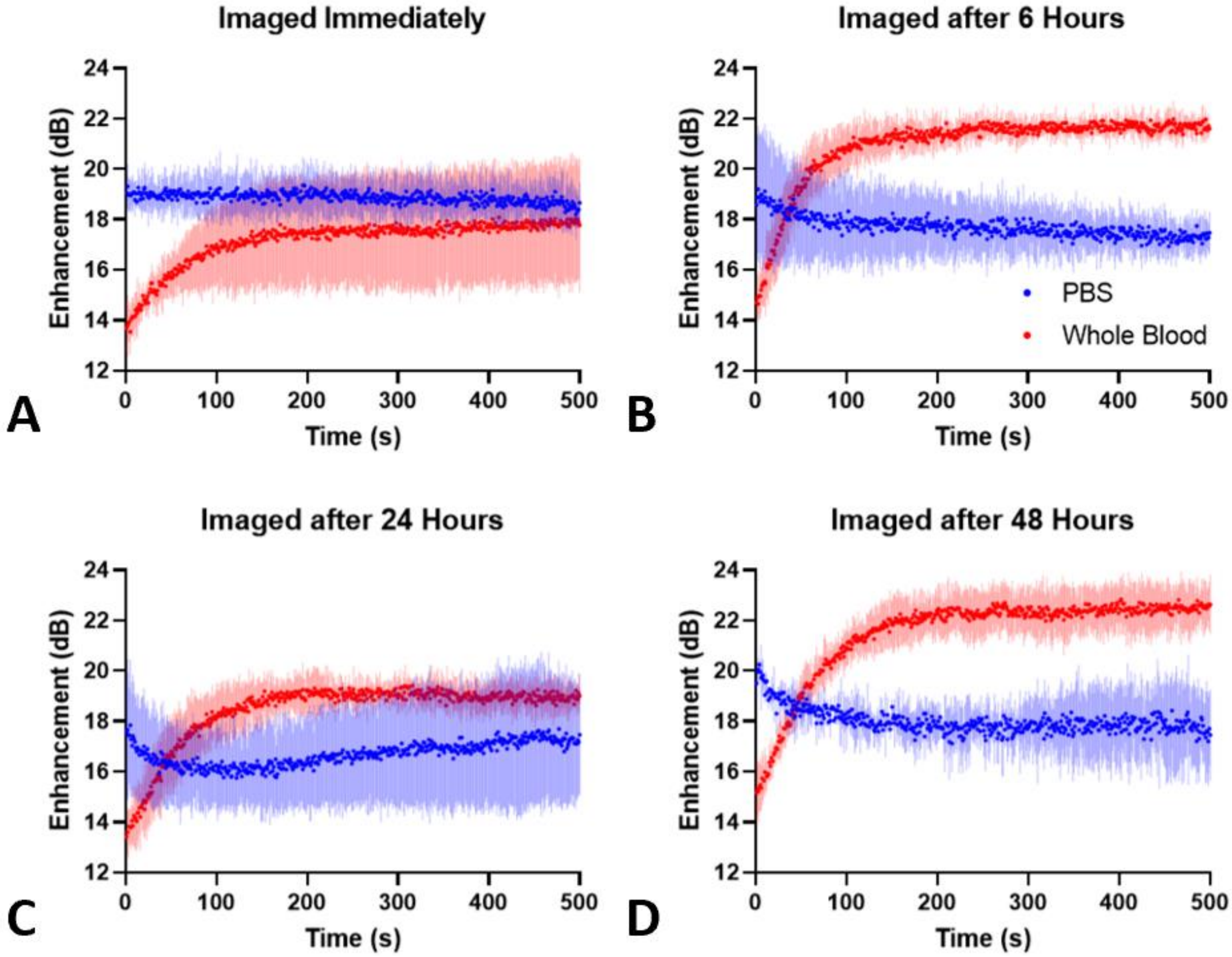


**Supplemental Figure 2. All experimental groups for whole blood vs PBS.** (a) Imaged immediately after mixing (b-d) Imaged at various time points post-initial mixing. Samples were stored at 4°C and imaged at 25°C. All whole blood groups show an increase in enhancement over time.

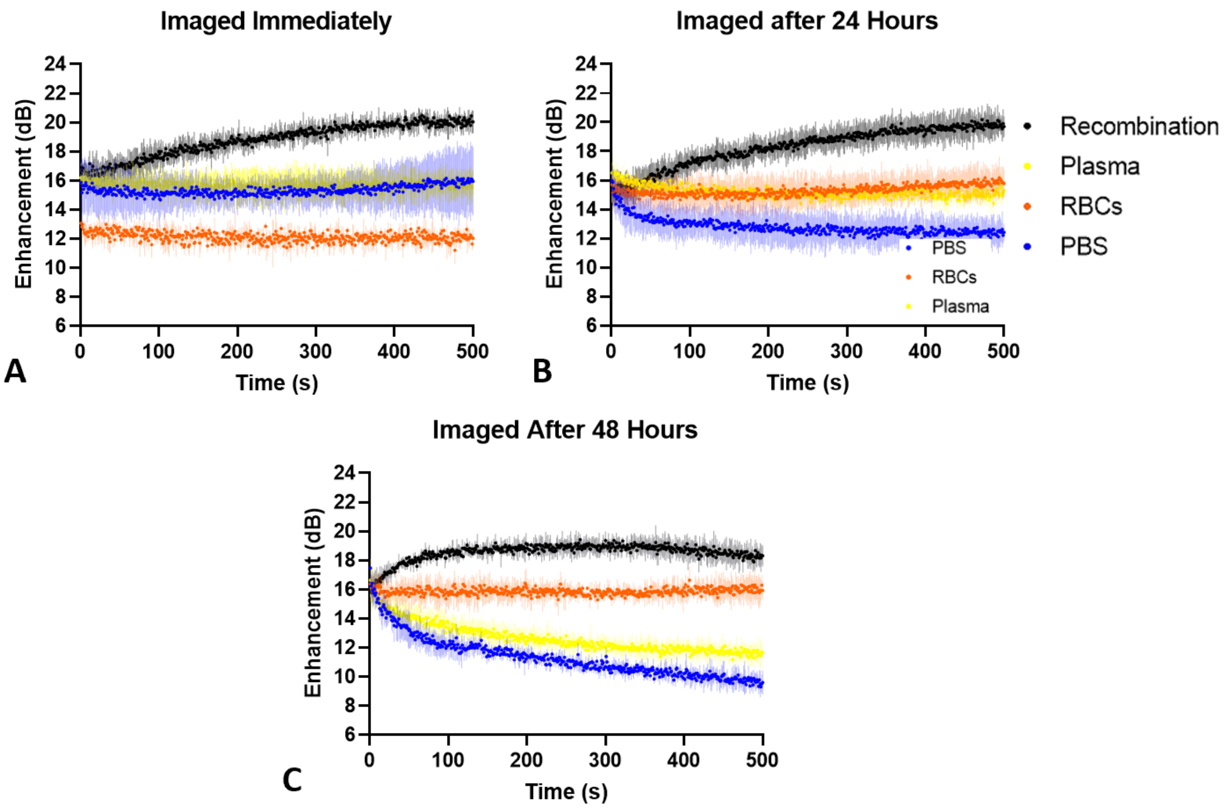


**Supplemental Figure 3. All experimental groups for RBCs, plasma, and PBS.** (a) Imaged immediately after mixing (b-c) Imaged at various time points post-initial mixing. Samples were stored at 4°C and imaged at 25°C. No groups show an increase in enhancement over time. The RBC group showed the most stability when imaged after 24 or 48 hours post initial mixing while PBS and plasma typically behaved similarly.

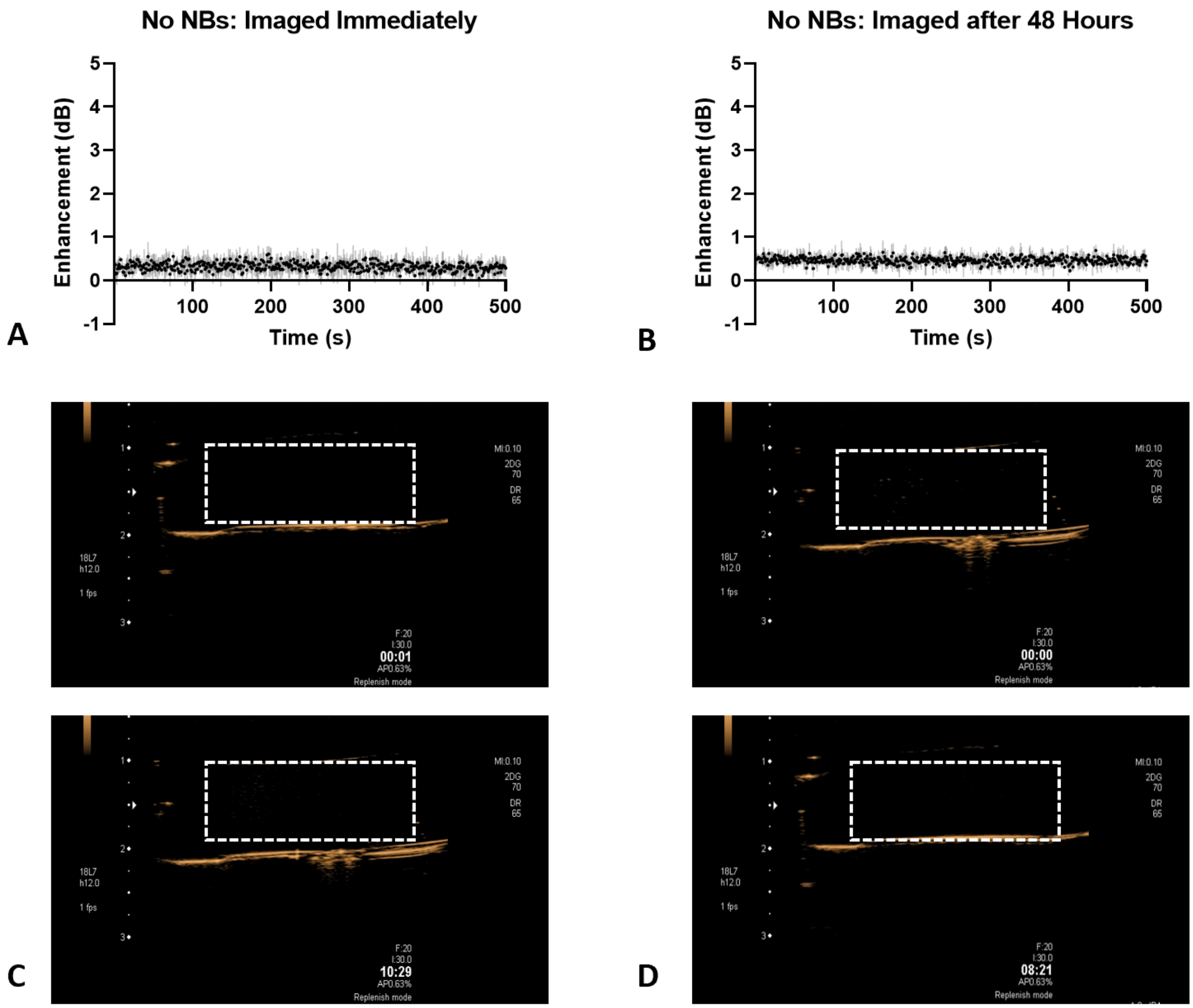


**Supplemental Figure 4. Enhancement over time of solution of whole blood with no nanobubbles.** (a-b) Solutions show minimal enhancement when imaged immediately and after 48 hours post initial mixing. (c) Visual representation of sample imaged immediately after mixing at 0 and 500 seconds. (d) Visual representation sample imaged 48 hours post initial mixing at 0 and 500 seconds. The white box represents the ROI and the phantom inlet where the blood solution is injected.

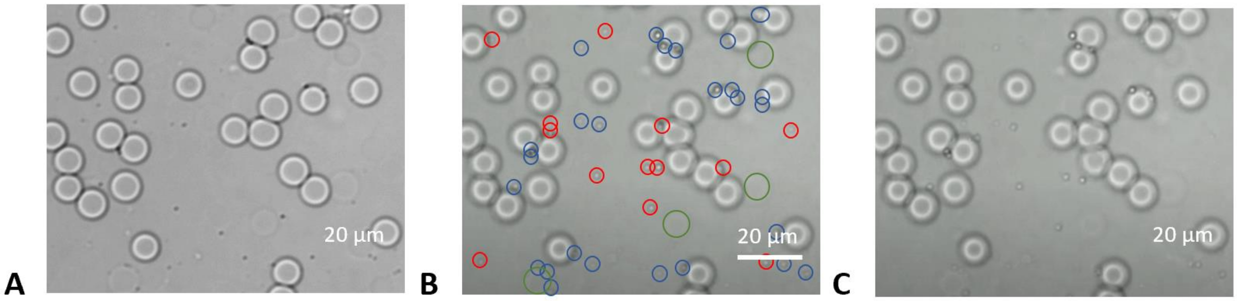


**D** See attached video submission

**Supplemental Figure 5. Light microscopy of NBs in whole blood.** (a) 63x in-focus images, (b) out-of-focus images at t=0 s, (c) out of focus images at t=4 s. Images are taken out of focus to emphasize NB location and verify that the particles are NBs. Red circles indicate NBs that moved location from t=0 to t=4 s and blue circles indicate NBs that did not move location. Larger green circles show RBCs that are out of plane. (d) 4 second video of NBs in whole blood at 63x in brightfield.

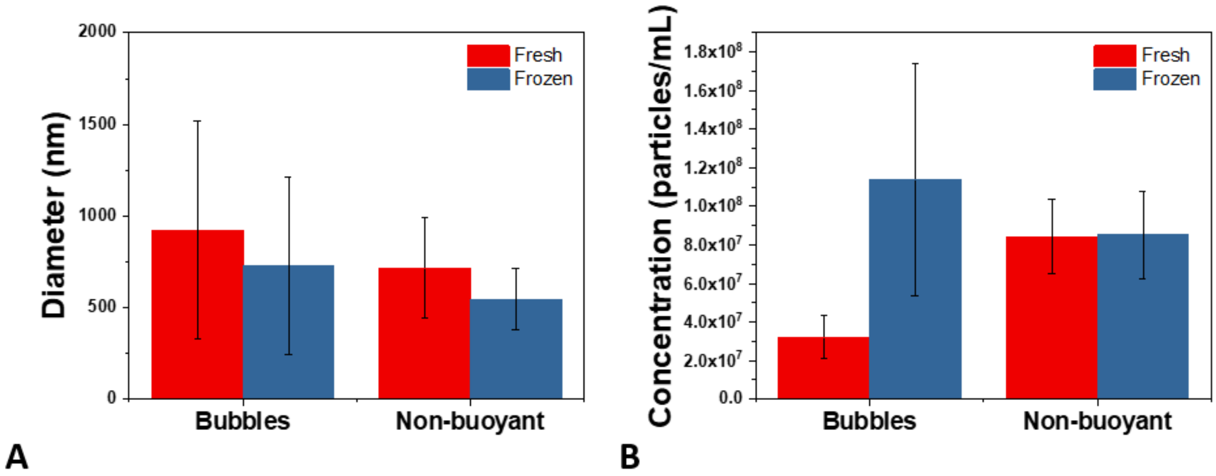


**Supplemental Figure 6. Lumason concentration and size analysis comparing fresh and frozen bubbles.** (a) Diameter of Lumason when the bubbles are used directly after activating (fresh) or frozen after activating and then thawed before use (b) Concentration of buoyant and non-buoyant Lumason when the bubbles are used fresh or frozen after activating. Bars represent standard deviation.
